# Evaluation of an optimized protocol and Illumina ARTIC V4 primer pool for sequencing of SARS-CoV-2 using COVIDSeq™ and DRAGEN™ COVID Lineage App workflow

**DOI:** 10.1101/2022.01.07.475443

**Authors:** Cyndi R. Clark, Matthew T. Hardison, Holly N. Houdeshell, Alec C. Vest, Darcy A. Whitlock, Dylan D. Skola, Jeffrey S. Koble, Michael Oberholzer, Gary P. Schroth

## Abstract

Next-Generation Sequencing based genomic surveillance has been widely implemented for identification and tracking of emerging SARS-CoV-2 variants to guide the Public Health response to the COVID-19 pandemic. Amplicon-based assays, such as the Illumina^®^ COVIDSeq™ Test (RUO) and COVIDSeq Assay (RUO), enable scalable sequencing of SARS-CoV-2, leveraging V3 and V4 primer designs from the ARTIC community and DRAGEN™ COVID Lineage App analysis available on Illumina BaseSpace™. We report here a comparison of COVIDSeq performance for SARS-CoV-2 genome reporting using the ARTIC V3 based primer pool (including primers for human control genes) that is provided with the COVIDSeq kit versus the ARTIC V4 based Illumina COVIDSeq V4 primer pool, using an optimized protocol and DRAGEN COVID Lineage App analysis. The data indicates that both primer pools enable robust reporting of SARS-CoV-2 variants. The Illumina COVIDSeq V4 primer pool has superior performance for SARS-CoV-2 genome reporting, particularly in samples with low virus load, and is therefore the recommended primer pool for genomic surveillance of SARS-CoV-2 for research use using COVIDSeq.

## Introduction

The first complete genome sequence of severe acute respiratory syndrome coronavirus 2 (SARS-CoV-2), the virus responsible for the global COVID-19 pandemic, was published worldwide within weeks of preliminary identification of the disease and launched the development of diagnostic tools and mRNA-based vaccines (ref 1). Since then, next-generation sequencing (NGS) of the viral genome has proven to be an indispensable tool for the detection and molecular epidemiology of SARS-CoV-2 variants and the introduction and spread of novel lineages with worsened pathological phenotypes. At the time of writing, close to one million SARS-CoV-2 sequences with metadata have been submitted to the GISAID database and SRA from NCBI (ref 2), representing the largest global genomic response to an individual infectious disease in history. This undertaking has emphasized the extent to which cost-effective NGS, optimized through standardized hybrid capture and amplicon-based workflows with ‘push-button’ analysis tools, has established itself as a critical tool for the Public Health response during current and future infectious disease outbreaks.

SARS-CoV-2 genomes accumulate ~2 point mutations per month (ref 3), primarily through errors during viral RNA replication. These mutations are used to partition sequences into phylogenetic trees that can identify disease outbreaks with a common origin and trace the spread of the virus around the world. When mutations are non-silent and occur in open reading frames (ORFs), they can have biologically relevant impacts on transmissibility and virulence, as well as reduced susceptibility to vaccine induced immunity and potential drug resistance. This is particularly the case in the gene encoding for the viral Spike protein (S gene), which interacts with the ACE2 receptor on the cell surface of human epithelia to initiate infection. Additionally, the Spike protein interacts with components of the host immune system, including antibody and cellular immune responses raised against vaccines, thus monitoring S gene mutations is critical for post-market surveillance of vaccines and antibody therapies. Other viral genes, such as the RNA-dependent RNA polymerase RdRP and the viral proteases are expected to gain major interest in the coming years as the targets of nucleoside analogs and protease inhibitors already on the market and under development. Scaling the use of antiviral drugs targeting these viral proteins will require post-market surveillance of the SARS-CoV-2 genome to monitor for the evolution and spread of mutations leading to antiviral resistance.

Since publication of the SARS-CoV-2 genome, the ARTIC network (ref 4) has released several primer schemes for tiled amplicon sequencing of SARS-CoV-2-positive clinical samples. Version 3 of this scheme, with 98 overlapping amplicons, was designed to cover the SARS-CoV-2 sequences available at the beginning of the outbreak (March 2020) and has been widely implemented in commercial assays (ref to COVIDSeq™) and community protocols (ref 5). The commercially available ARTIC V3 based primer pools included in the Illumina COVIDSeq™ Test (RUO) and COVIDSeq™ Assay (96 samples) also include a set of primers for amplification of human mRNAs, which serve as internal controls for the assay.

Over the course of the pandemic, genetic variation has supported the evolution of a number of SARS-CoV-2 descendant lineages. Four variants (Alpha, Beta, Gamma, and Delta) have been the dominant sources of infection with the Delta Variant of Concern (VOC) accounting for the majority of new cases around the world as of November 2021 (GISAID). Some of these VOC mutations co-localize with the primer binding sites of the ARTIC V3 scheme, which can thermodynamically destabilize the primer-cDNA duplex required for PCR amplification causing reduced or missing read coverage for portions of the genome with these primer binding site mutations, especially in samples with low viral titer or degraded RNA. The Spike ORF has been particularly impacted by viral evolution, due to the exposure of the encoded Spike protein to the immune system of different hosts. Given the central importance of the Spike protein to both infection and immune responses, it is vital to monitor mutations in its coding gene. As a result, the Spike ORF contains a higher frequency of mutations, including primer binding site mutations, than other regions of the SARS-CoV-2 genome.

The ARTIC V3 primer design is considered the ‘work horse’ for SARS-CoV-2 amplification and subsequent NGS analysis and can achieve over 90% genome coverage in Variants of Concern. However, reports indicate that this primer pool may lead to lower coverage of certain amplicons, such as 72, 74, 76 (ref 6). This reduced performance impacts consensus sequence generation and high-resolution coverage of the spike protein gene in recent variants, such as Delta, particularly in low viral titer samples. Strategies to mitigate the impacts of primer binding site mutations include increasing the number of amplification cycles to ensure maximum amplification is reached for all amplicons, modified PCR thermal cycling protocols, and spike in primers to complement the ARTIC V3 primer set.

With an increasing need to update the ARTIC V3 primer design to evolve with the virus, the ARTIC community designed a novel primer scheme ARTIC V4, taking into consideration the genetic variability and high frequency mutations in the dominant variants circulating in June 2021 (B.1.1.7, B.1.351, B.1.429, B.1.525, B.1.617.1, B.1.617.2 and P.1.) (ref 6). The ARTIC V4 design produces 99 overlapping amplicons and has been optimized to amplify across the entire viral genome with the same efficiency through variable concentrations of the primer pairs (ref 6).

The Illumina COVIDSeq™ Test (RUO) is a 3072-sample kit that is widely used for surveillance of SARS-CoV-2 variants using NextSeq™ and NovaSeq™ 6000 in combination with the Illumina DRAGEN™ COVID Lineage App for ‘push-button’ variant calling and mutation reporting through Illumina BaseSpace Sequence Hub. The recently launched COVIDSeq™ Assay is a 96-sample kit, ideally suited for de-centralized surveillance using small batches of samples on Illumina iSeq100™, MiniSeq™ and MiSeq™ systems. Both commercial products are supported with a primer pool based on the ARTIC V3 and primer combinations targeting a set of human mRNA controls. Recently, Illumina commercialized a novel, stand-alone primer pool, based on the ARTIC V4 design (Illumina^®^ COVIDSeq™ V4 Primer Pool, cat number 20065135). The Illumina DRAGEN™ COVID Lineage App available on Base Space Sequence hub has been updated to allow the selection of ARTIC V3 or ARTIC V4 primer schemes for analysis and variant calling, depending on the primer pool used (ref).

In this study we evaluated the performance of Illumina COVIDSeq™ Test (RUO) in combination with the Illumina^®^ COVIDSeq™ V4 Primer Pool and the DRAGEN™ COVID Lineage App for whole genome sequencing, alignment, and variant calling of recently identified SARS-CoV-2 positive specimens with a wide range of Ct values.

## Methods

### Sample Preparation

Upper respiratory tract specimens were collected from individuals experiencing COVID-19-related symptoms in a variety of clinical settings using a flocked nasal swab and Longhorn PrimeStore^®^ Molecular Transport Media (MTM). The King Fisher™ Flex Magnetic Particle Processor with 96 Deep-Well head and the MagMAX™ Viral/Pathogen II Nucleic Acid Isolation Kit was used for high throughput extraction of patient specimens, and the TaqPath™ COVID-19 Combo Kit was used for the detection of SARS-CoV-2 ORF1ab, N protein, and S protein genes by real-time RT-PCR using Applied Biosystems^®^ QuantStudio™ 7 Flex Real Time PCR instruments, QuantStudio™ Real Time PCR Software v1.3, and Applied Biosystems^®^ COVID-19 Interpretive Software.

Specimens with an average cycle threshold (Ct) value ≤ 35 were selected from a broad geographic region and sorted based on a Ct < 30 or Ct of 30-35. A total of 1504 positive specimens were selected for re-extraction, library prep, and sequencing, with 335 samples having a Ct value <30 and 1169 samples having a Ct value in the range of 30-35.

A 400 μl aliquot of each positive specimen selected for NGS was extracted and eluted in 50 μl of nuclease-free water. Library preparation was automated via Tecan EVO liquid handlers following the Illumina COVIDSeq™ RUO Reference Guide (# 1000000126053) workflow with the following modifications:

1. All steps were performed in 384-well PCR plates including 376 positive specimens, 4 positive controls, and 4 negative controls from 4, 96-well extraction plates
2. ‘Amplify cDNA’ PCR program was changed to include 10 additional PCR cycles to increase the amplicon coverage and improve sensitivity of the assay
  a. 98°C for 3 minutes
  b. 45 cycles of:
    i. 98°C for 15 seconds
    ii. 63°C for 5 minutes
    iii. Hold at 4°C
3. ‘Amplify cDNA’ COVIDSEQ™ PCR Master Mix 1 and 2 were formulated using either the V3 primer pools (provided in the COVIDSEQ™ Library Preparation kit) or an aliquot of the V4 primer pools manufactured by Illumina, Inc for direct comparison of all specimens.
4. ‘Tagment PCR Amplicons’ reaction volumes were reduced to accommodate the 384-well PCR plate.
  a. 5 μl cDNA 1
  b. 5 μl cDNA 2
  c. 15 μl Master Mix
5. ‘Post Tagmentation Clean Up’ was skipped to avoid losing amplicons.
6. ‘Amplify Tagmented Amplicons’ reaction volumes were reduced to accommodate the 384-well PCR plate.
  a. Amplicons bound to magnetic beads without supernatant
  b. 20 μl Master Mix
  c. 10 μl indexes

Tagmented amplicons were pooled, normalized, and diluted, and 0.4 nM libraries were sequenced on the NovaSeq™ 6000 at 2 x 150 cycling using a S4 Flowcell, conditions for a targeted sequencing depth of 1M read-pairs (2M paired-end reads) and uploaded to the Illumina BaseSpace™ Sequence Hub (BSSH) cloud analysis service.

### Data Analysis

FASTQ files were generated from the Run data using the FASTQ Generation v1.0.0 BSSH app. These files were analyzed using the DRAGEN™ COVID Lineage app (using a pre-release version v3.5.613 that supported both V3 and V4 primers) with the following Advanced Settings (General) for both primer schemes.

‘Enable ZIP of most output files’ unchecked
‘Aligner Min Score’ changed to 8
‘Consensus Sequence Generation Threshold’ changed to 5

This software performs a k-mer based detection step that tests for the presence of a subset of tiled 32-mers within the Wuhan reference genome across all reads in the sample. If at least 150 k-mers for a given amplicon are detected at least once, that amplicon is considered detected. If at least five amplicons are detected in this way, the sample is considered SARS-CoV-2 positive, and mapping with the DRAGEN™ read aligner is performed, followed by variant calling using the DRAGEN™ somatic variant caller. High quality (PASS value in filter column, minimum DP 10, minimum AF 0.5) variants are then passed to a consensus sequence module that masks any genomic regions with less than 10X coverage and uses bcftools (ref 9) to generate a consensus viral genome sequence, based on the NC_045512.2 reference genome that contains these high-quality variants.

The outputs of DRAGEN™ COVID Lineage were remotely accessed using the Basemount software package and Aegis-developed BaseSpace™ CLI Download Assistant application to query the projects in Illumina’s BaseSpace™ Sequence Hub using command line interface to download consensus metrics and coverage metrics for each sample. Per-bp read coverage (after removing duplicate reads) across the viral genome was computed from the files output by the DRAGEN™ aligner. Since both the V3 and V4 primer sets generate overlapping amplicons, the mean coverage in each amplicon was calculated over the non-overlapping portion of each amplicon. The mean amplicon coverage values were then transformed by taking the base-10 logarithm after adding a pseudocount of 1.

Variant call quality was assessed using the Nirvana software package to annotate the set of high-quality variants called for each sample. Any variant predicted to cause the gain or loss of a translation initiation or termination site is unlikely to be biologically feasible, so the frequency of such variant calls can be used as a proxy for the frequency of false-positive variant calls. Although this method is expected to miss many true false-positive calls in the absolute sense, using it for differential comparisons, as we do here, avoids this pitfall. To test whether the rate of putative false positive variant calls per sample differed between the V3 and V4 primers, we constructed nested maximum likelihood Poisson models of the number of putative FP variants per sample. The likelihood ratio test was employed to compare the model wherein the data for both primer sets were generated with the same mean to one in which the data were generated with different means for the V3 and V4 primers. To test for differences in the rate of putative false positives per variant call, we applied the same procedure but substituted Bernoulli models in place of Poisson models.

## Results

Completed sequencing data for 1315 samples processed with both ARTIC V3 and V4 primer schemes over a range of RT-PCR Ct values was analyzed, with most of the samples having an average Ct value > 30 (Figure 1A). To avoid sequencing low-quality data, samples with less than 5 detected amplicons are not processed through alignment and variant calling. 1269 samples produced detectable sequences (Figure 1B). 308 samples were detectable with only the V4 primer set than versus 131 samples detectable only with the V3 primer set (Figure 1B). 830 samples produced a detectable sequence with each primer set V3 and V4 and were included in further analysis.

**Figure 1:**
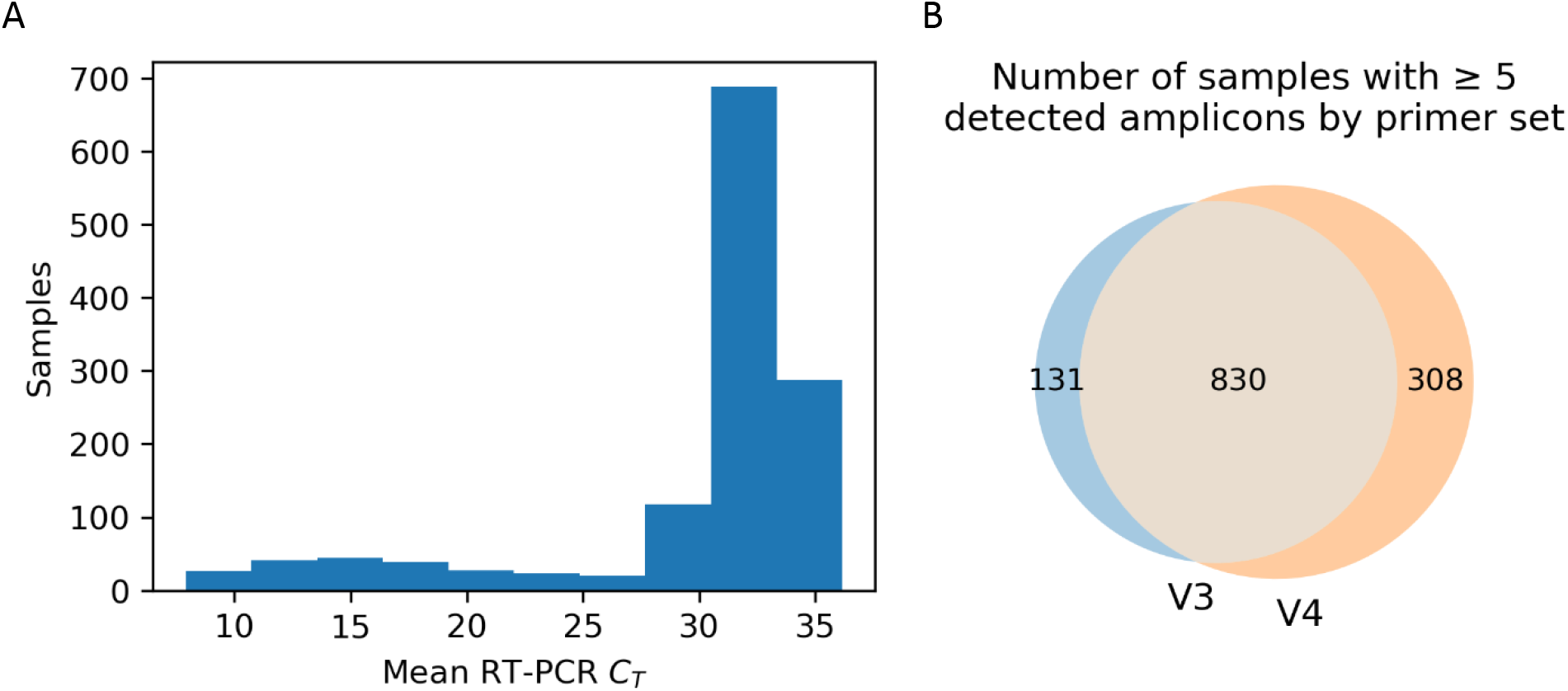
(A) Histogram of mean of RT-PCR CT values against 3 targets (Orf1a, N gene, and S gene) for samples in this study. (B) Number of samples with ≥ 5 detected amplicons by primer set. Read alignment and variant calling was not performed on samples with less than 5 detected amplicons. Of the cohort of 1315 samples, 46 did not produce ≥ 5 detectable amplicons with either primer set.

To evaluate the performance of V3 versus V4 primer pools, we first compared the number of callable bases over each amplicon covering the virus genome (Figure 2A). The results indicate a substantial reduction in non-callable bases ‘N’ when using the V4 primers, in comparison to V3 primers, particularly in ≥ 30 Ct (low virus titer) samples (Figure 2A). Importantly, the non-callable bases ‘N’ value is substantially reduced in the Spike gene region when using V4 primers (Figure 2A). As can be seen in figure 2B, we observed a significant increase in the number of detected amplicons in both the low CT samples (mean 91.17 detected amplicons for V3 primers, 95.37 for V4 primers, p=1.37E-36 by one-sided Wilcoxon signed rank test) and high CT samples (mean 19.94 detected amplicons for V3 primers, 30.77 for V4 primers, p=1.00E-74 by one-sided Wilcoxon signed rank test). The same pattern was observed for median non-duplicate read coverage across both low CT (mean 5680.22 median coverage for V3, 6602.94 for V4, p=0.023 by one-sided Wilcoxon signed rank test) and high CT (mean 59.12 for V3, 167.58 for V4, p=1.87×10E-79 by one-sided Wilcoxon signed rank test) sample cohorts. The smaller difference in median coverage in the low CT samples is likely a consequence of saturating coverage in high titer samples due to PCR exhaustion and duplicate removal.

**Figure 2:**
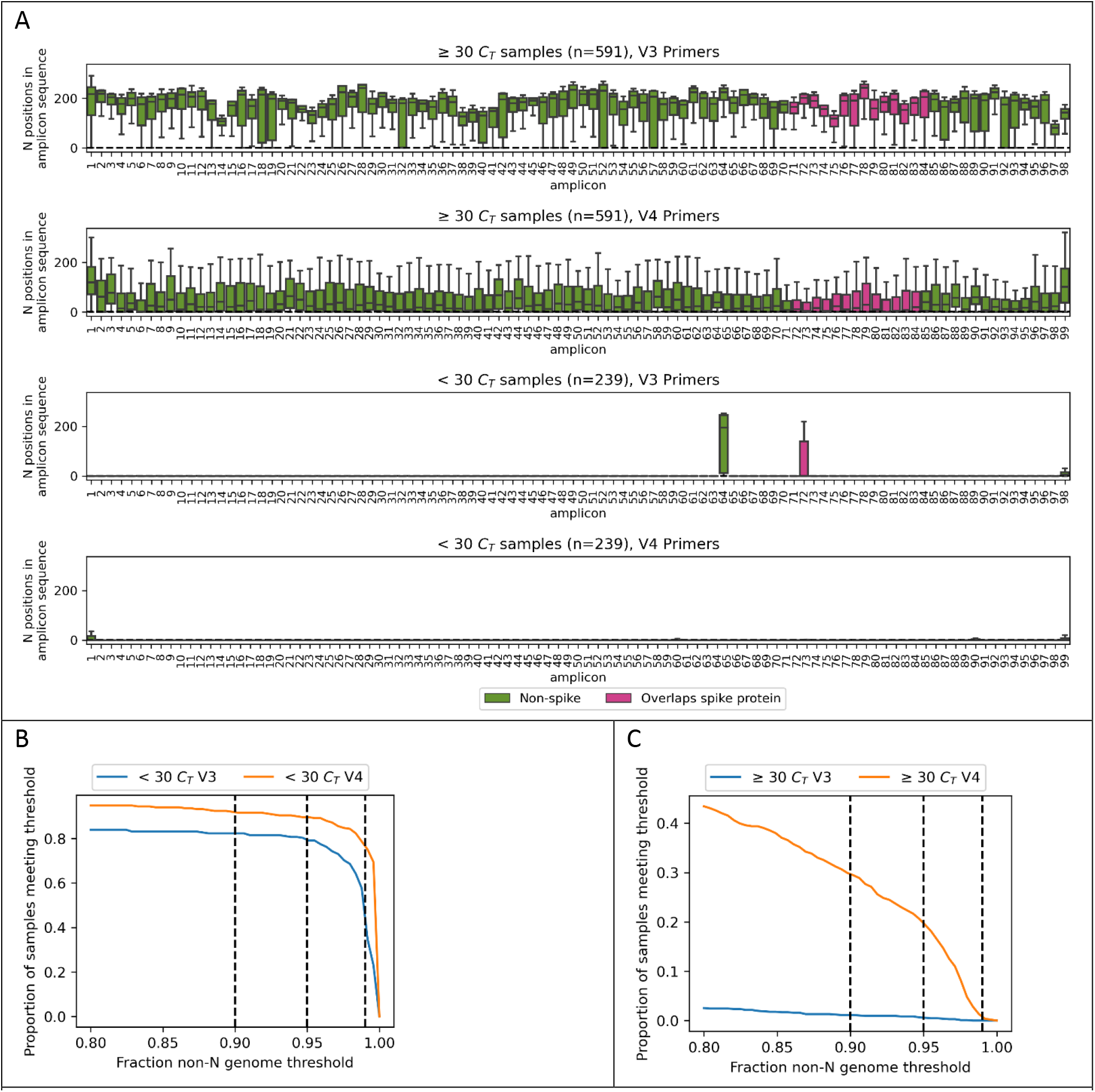
(A) Boxplot of number of Ns in each amplicon sequence for samples with at least 5 detected amplicons in both primer sets, partitioned by mean CT and primer set. Amplicons that overlap the Spike protein CDS are colored magenta. (B) Detail of cumulative distribution of fraction non-N genome values for samples with mean CT < 30. Dashed vertical lines indicate commonly used quality thresholds of 0.90, 0.95, and 0.99. (C) Same as (B) for samples with mean CT >= 30.

The fraction of the consensus viral genome that is not “N” is an important metric for the utility of viral surveillance methods, since it indicates how much of the viral genome could be reliably sequenced. Best practices in viral resequencing dictate that, in order to avoid reference bias, genomic positions in which the allele could not be determined should be hard-masked with the “N” character (for the DRAGEN™ COVID Lineage software used in this analysis, positions with less than 10X read coverage are masked). Most sequence repositories have minimum fraction-non-N requirements for accepting or marking viral genomes as high-quality (Ref 2). As can be seen in cumulative distribution of fraction non-N values for low CT samples in Figure 2C, the V4 primer set generates nearly twice (191 to 107) as many sequences passing the 0.99 fraction non-N threshold, with even more pronounced improvement and in the high CT samples, with 319 having at least 0.90 non-N content with the V4 primers compared to 12 for the V3 primers (Figure 2D)

To ensure that the V4 primer set was not negatively impacting the accuracy of variant calls, we performed an estimation of relative precision between the two primer sets. We used Nirvana to annotate the likely consequences of each variant in the 830 samples with at least 5 detected amplicons with both primer sets. We counted any variant that caused the gain or loss of either a translation start or stop site to be a “putative false-positive” since such events are likely to be highly deleterious to viral fitness and are therefore expected to be strongly selected against in the population of true sequence variants. Across all samples, we found 32 out of 14378 putative false positive variant calls using the V3 primers, and 43 out of 16917 calls using the V4 primers. Two-sided tests for the V3 and V4 primers having different rates of putative false positive variants per sample and per variant call were not significant (table 1).

**Table 1:**
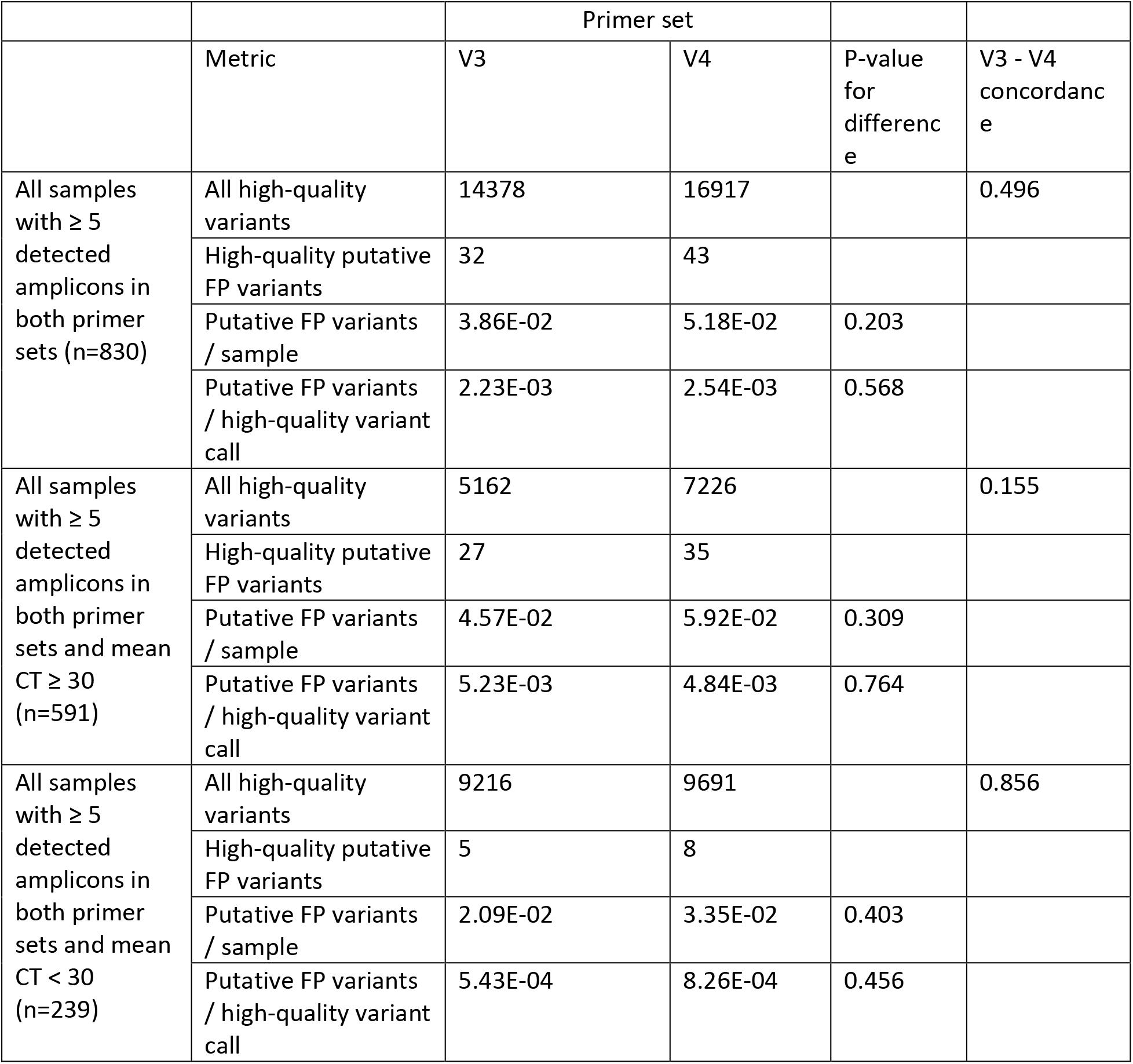
Counts of all high-quality variants and putative false positive (variants that introduce or remove a translation start or stop site) high-quality variants for all samples that had at least 5 detected amplicons in both samples (n=830) as well as for the subsets of samples at or above 30 CT and below 30 CT. P-values are given for a likelihood ratio test for the improvement given by using separate Poisson (for FP counts by sample) or Bernoulli (FP counts per variant) models for the V3 and V4 primer sets compared to a single model for both primer sets.

## Discussion

Sequencing of SARS-CoV-2 novel variants, including Delta, with high accuracy is a critical Public Health need. Although the SARS-CoV-2 Delta variant is currently dominant worldwide, the virus continues to evolve and Delta sub-variants are emerging, requiring continuous update of amplicon-based primer pools to scope with virus mutations. Assessing the performance of workflows on large number of representative recent clinical samples across relevant viral titers is critical to provide insight into the performance of SARS-CoV-2 sequencing assays. To our knowledge, this is the first published, large-scale, experimental study comparing ARTIC V3 vs ARTIC V4 primer designs on recent clinical remnant samples using commercially available assays and analysis tools.

From 1315 samples included in the study, 830 samples generated a sequence with ≥5 amplicons detected by the V3 and the V4 primer design, and these samples were included in further analysis. From the 830 samples, 591 samples had Ct value of ≥30 and 239 samples had Ct value <30, respectively, representing relevant biological range as expected in samples generally used for reporting SARS-CoV-2 virus sequences.

We first compared the number of non-callable bases (N) in sequences generated with primer design V3 versus V4, which is a critical value for reporting a virus sequence to public databases (ref 7). In the 591 samples with Ct value ≥30, representing moderate to low viral titer samples, the V4 primer set was able to report more callable bases in each amplicon, as indicated by a decrease in N-positions in amplicons covering the virus genome. To ensure high quality data, the global databases, such as GISAID, require a 90% of breadth of coverage over the virus genome to report a sequence (ref 7)). V4 primers generates approximately twice as many sequences passing the 90%, 95% and 99% fraction non-N thresholds enabling an overall higher number of samples to be reported to global databases using V4 primers in comparison to V3 primers. This difference is particularly evident in ≥30 Ct (low viral titer) samples, which represent many routine samples obtained by laboratories as diagnostic remnants for NGS sequencing follow up. The difference between V3 and V4 primers was also evident in the sample set <30 Ct (high viral titer), however the difference was less pronounced than in the ≥30 Ct (low viral titer) sample set. Therefore, the V4 primers provide a substantial advancement versus V3 primers for routine surveillance of SARS-CoV-2 using COVIDSeq™, particularly in low-to-moderate viral titer samples, as commonly used in the ongoing surveillance effort.

In addition to maximizing the number of samples that can be reported to global surveillance repositories, sequence coverage in the Spike gene is of particular relevance, due to the increasing number of mutations in this gene and their potential biological relevance. Reports indicate a decreased coverage of successful viral sequencing in the Spike gene when using V3 primer designs in amplicon-based workflows for SARS-CoV-2 sequencing (ref 6)), most likely due to viral evolution. The number of N-positions (non-callable bases) in amplicons overlapping the Spike gene is significantly reduced with the V4 primers versus the V3 primers in ≥30 Ct samples (mean 110.02 N positions per Spike amplicon for V3 primers, 37.83 for V4 primers, p=8.93E-89 by one-sided Wilcoxon signed rank test), indicating that the V4 primer design provides improved sequence of the Spike gene in recent variants, including amplicon regions know to be problematic with V3 design (72, 76, 78). In < 30 Ct (high virus titer) samples, the V3 primers provide a comprehensive coverage of samples analyzed, with the exception of amplicons 64 and 72. Conversely, the V4 primers enable comprehensive coverage of the viral genome, including the V3 amplicon 64 and 72 regions, providing a critical improvement in sequencing the Spike gene in currently circulating variants (mean 14.27 N positions per Spike amplicon for V3 primers, 4.37 for V4 primers, p=1.61E-13 by one-sided Wilcoxon signed rank test),.

Some of this improvement in COVIDSeq™ performance is expected to be the result of the V4 primer binding sites designed to avoid regions with mutations in the currently circulating variants, particularly for the problematic genomic regions covered by amplicons 72 and 64 in the V3 scheme. Another likely advantage of the V4 primer pool is the removal of the human control primers, thus allowing the reagents used in the library preparation steps to solely amplify viral material.

Amplicon-based approaches, such as COVIDSeq™, provide a streamlined and scalable NGS workflow at low cost, enabling a high-throughput centralized and low-throughput de-centralized surveillance response to the rapidly evolving SARS-CoV-2 virus. Amplicon approaches depend on simultaneous and efficient annealing of a large number of primers to their viral target regions, which is impacted by mutations in the primer target regions. Therefore, amplicon-based approaches depend on continuous updates to primer combinations and bioinformatic tools that can function, irrespective of virus evolution and to accommodate mutations in primer binding sites. The transition from the ARTIC V3 to ARTIC V4 primer set is an example of how amplicon-based sequencing can be maintained at high efficiency for reporting the genome of a rapidly evolving virus, wherein real-time updating of primer combinations for established workflows and analysis tools is critical.

Although the V4 primer set has improved the SARS-CoV-2 Delta strain genomic coverage, there is a possibility that a new design will be required in the future, or even reversion to the V3 set as more strains emerge. It is critical to establish standardized workflows for NGS library preparation, along with bioinformatic tools, to enable the integration of novel primer combinations in response to viral mutation. COVIDSeq™ in the 3072 and 96 sample kit formats, in combination with the ‘push-button’ COVID Lineage App provides these building blocks, enabling the use of various primer combination such as ARTIC V3 and V4 primers, as well as future primer combinations. The analytical processes created by COVIDSeq™ and the DRAGEN™ COVID Lineage App, and strategies for updating primer combinations, as led by the ARTIC community, provide a critical blueprint for surveillance solutions to emerging and re-emerging viruses and Antimicrobial Resistance with pandemic potential.

